# Characterisation of *Ornithobacterium hominis* colonisation dynamics and interaction with the nasopharyngeal microbiome in a South African birth cohort

**DOI:** 10.1101/2025.05.24.655922

**Authors:** Celine C De Allende, Susannah J Salter, Siobhan E Brigg, Shantelle Claassen-Weitz, Kilaza S Mwaikono, Lesley Workman, Heather J Zar, Mark P Nicol, Julian Parkhill, Felix S Dube

## Abstract

*Ornithobacterium hominis* is a recently described Gram-negative bacterium that colonises the human nasopharynx and may be associated with poor upper respiratory tract health. Here, we describe the isolation of *O. hominis* from samples collected from a South African birth cohort, creating the first archive of cultured strains of the species from Africa. Sequenced genomes from this archive reveal that South African *O. hominis* is more similar to Australian strains than those from Southeast Asia, and that it may share genes with other members of the microbiome that are relevant for virulence, colonisation, and antibiotic resistance. Leveraging existing microbiome data from the cohort, *O. hominis* was found to be closely associated with bacterial co-colonisers that are rare in non-carrier individuals, including *Suttonella*, *Helcococcus*, *Moraxella* spp., and Gracilibacteria. Their collective acquisition has a significant impact on the diversity of nasopharyngeal communities that contain *O. hominis*. Individuals who have not yet acquired *O. hominis* have a higher abundance of *Moraxella* (particularly *M. lincolnii*) than individuals who never acquire *O. hominis*, suggesting that this could be a precursor state for successful colonisation. Finally, a novel co-coloniser species, *Helcococcus ekapensis*, was successfully isolated and sequenced.

**Data Summary:** *Ornithobacterium hominis* data have been deposited under project accession ERP149886. This comprises genome sequences for isolates SA-OH-C1 (ERR13967269), SA-OH-C2 (ERR13967270), SA-OH-C3 (ERR13967271), SA-OH-C4 (ERR13967272, ERR13967275), SA-OH-C5 (ERR13967273), SA-OH-C6 (ERR13967274, ERR13967276). Previously published 16S rRNA gene data are deposited under project accessions PRJNA790843 and PRJNA548658. *Helcococcus ekapensis* genome data are deposited under project accession PRJEB85661.

**Software used:**

- AMRFinderPlus v3.12.8: https://github.com/ncbi/amr
- AssembleBAC-ONT v1.1.1: https://github.com/avantonder/assembleBAC-ONT
- BAKTA v1.8.1: https://bakta.computational.bio/
- BLAST v2.16.0: https://blast.ncbi.nlm.nih.gov/Blast.cgi
- Comprehensive Antibiotic Resistance Database (CARD) Resistance Gene Identifier (RGI) tool v1.2.1: https://card.mcmaster.ca/analyze/rgi
- Decontam v1.12 (R package): https://github.com/benjjneb/decontam
- Eggnog-mapper v2.0.1: http://eggnog-mapper.embl.de/
- FastANI v1.1.0: https://github.com/ParBLiSS/FastANI
- Flye v2.9.2: https://github.com/fenderglass/Flye
- Guppy v6.5.7: https://community.nanoporetech.com/downloads/guppy/release_notes
- ISEScan v1.7.2.3: https://usegalaxy.eu/root?tool_id=toolshed.g2.bx.psu.edu/repos/iuc/isescan/isescan/1.7.2.3+galaxy1
- Medaka v1.9.1: https://github.com/nanoporetech/medaka
- MEGA11: https://www.megasoftware.net/
- MMseqs2 v17: https://github.com/soedinglab/MMseqs2
- Mothur v1.44.3: https://github.com/mothur/mothur
- NetCoMi v1.2.0 (R package): https://github.com/stefpeschel/NetCoMi
- Panaroo v1.4.3: https://github.com/gtonkinhill/panaroo
- PHASTEST: https://phastest.ca/submissions/new
- Prowler (commit ID c3041ba): https://github.com/ProwlerForNanopore/ProwlerTrimmer
- R v4.4.3: https://www.r-project.org/

**Databases used:**

- Comprehensive Antibiotic Resistance Database (CARD): https://card.mcmaster.ca/
- European Nucleotide Archive: https://www.ebi.ac.uk/ena/
- Genome Taxonomy Database (GTDB) release 09-RS220: https://gtdb.ecogenomic.org/
- RefSeq release 228: https://www.ncbi.nlm.nih.gov/refseq/about/prokaryotes/
- SILVA v132: https://www.arb-silva.de/

**Impact statement:** First described in 2019, *Ornithobacterium hominis* is an understudied bacterium that may be associated with poor respiratory health in children. The study builds upon existing knowledge of *O. hominis* by describing the first African isolates of the species, its potential as a reservoir of virulence and antibiotic resistance genes in the upper respiratory tract, and the unique microbiome profile of *O. hominis* carriers.

## Introduction

Lower respiratory tract infections are a major cause of morbidity and mortality in children worldwide, especially in low- and middle-income countries (LMICs) (1). For example, lower respiratory tract infections caused by respiratory syncytial virus lead to millions of hospitalisations globally each year, but 99% of the resulting deaths in children under five occur in LMIC settings (2). The leading cause of bacterial pneumonia, *Streptococcus pneumoniae* infection, caused over 340,000 deaths in children under five in 2016, half of which were in sub-Saharan Africa (3).

Invasive bacterial infection is usually preceded by colonisation of the upper respiratory tract including the nasopharynx (4) and vaccination and widespread use of antibiotics may perturb the colonisation dynamics of these microbes (5). The nasopharynx harbours a diverse community of microorganisms. Some nasopharyngeal pathogens have been shown to negatively impact one another’s growth, for example the inverse correlation of *S. pneumoniae* and *Staphylococcus aureus* carriage due to pneumococcal H2O2 production and hydroxyl radical conversion (6,7). The “health associated” bacterium *Dolosigranulum pigrum* can also inhibit the growth of *S. aureus* and, in combination with *Corynebacterium*, inhibit the growth of pneumococci (8). Conversely, bacteria may rely on one another for nutrients to support growth, such as *Haemophilus influenzae* benefitting from factor V (NAD) released by the haemolytic activity of *S. aureus* (9).

*Ornithobacterium hominis* is a Gram-negative bacterium that can colonise the human nasopharynx persistently and at high abundance (10). Genomes from this species were first described from metagenomic samples from Thailand in 2019 (11) and from the only cultured isolates thus far, from Australia (12)*. O. hominis* has since been identified in microbiome data from countries across the world including South Africa (13), and carriage may be associated with poor upper respiratory tract health such as purulent rhinorrhoea and recurrent otitis media (14). However, little is known about the organism’s interactions with other members of the microbiome or its potential as a reservoir of antibiotic resistance or virulence factors.

We aimed to investigate the genetic diversity and colonisation dynamics of *O. hominis* in a South African birth cohort, and to describe the differential microbiome profiles of children colonised with *O. hominis* compared to uncolonised counterparts.

## Methods

### Sampling and participants

The Drakenstein Child Health Study (DCHS) (15) is a population-based birth cohort in which pregnant mothers in Paarl, South Africa, were enrolled during their second trimester and mother– child pairs have been followed from birth until at least when the children reach adolescence.

Nasopharyngeal swabs were collected every two weeks from birth through to one year of age in an intensive cohort or at well baby visits until 12 months of age, and every six months thereafter.

Archived nasopharyngeal samples were stored at –80 °C. Ethical approval was received from the Human Research Ethics Committee of the University of Cape Town, South Africa (401/2009 and 585/2015).

### 16S rRNA gene data and screening

Data from a prior DCHS microbiome study (13) was used to identify *O. hominis* enriched samples: 6,025 nasopharyngeal specimens and 330 induced sputum samples from 608 infants were examined through 16S rRNA gene sequencing targeting the V4 region, along with 754 technical controls. Detailed library preparation and sequencing methods have been described previously (16).

To enable rapid identification of *O. hominis* in a large set of samples, the amplicon dataset was screened for perfect matches to three sequences unique to this species in the 16S V4 region: [GAGCGTTATCCGGATTCATTGGGTTTAAAGGGTCYGTAGGCGGGCTRATAAGTCAGTGGTGAA ATCTCAC], [GAGTGAGTTTGATGTTGCTGGAATGTGTAGTGTAGCGGTG], and [ATGCGAAGGCAGGTAACAAAGACTTAACTG].

### *O. hominis* isolation

Based on 16S rRNA gene abundance data and prior metagenomic data, 36 nasopharyngeal samples with >1% relative abundance of *O. hominis* reads were identified for isolation attempts, giving rise to six isolates. Nasopharyngeal swabs were inoculated onto Columbia agar plates supplemented with 5% sheep’s blood (CBA) and incubated at 30 °C for at least 120 hours under microaerophilic conditions. Oxoid™ CampyGen™ 2.5 L sachets (ThermoFisher Scientific) were used to generate a microaerophilic environment. Presumptive *O. hominis* colonies, as described previously (12), were selected and sub-cultured on CBA and incubated at 30 °C for at least 48 hours. Colonies from primary culture appeared greyish, punctiform, and only apparent after five days of incubation. Colonies were sub-cultured until pure cultures were obtained.

Colonies were subjected to PCR targeting the conserved *O. hominis toxA* gene, using a procedure modified from that described previously (11). Briefly, a single colony was transferred to 200 µL of AVE buffer (Qiagen, Germany) using a sterile loop and heated at 95 °C for 5 minutes. The lysate was diluted 1:10 in nuclease-free water and PCRs were performed using 1 µL of diluted lysate in a final reaction volume of 25 µL with OneTaq® Quick-Load® 2× Master Mix (New England Biolabs) and OH_TOXIN primers (Inqaba biotech, South Africa) targeting the *toxA* gene: OH_TOXIN-F_ii 5′-GATGTATTGATAGATACTCCCGCCATTACG-3′, OH_TOXIN-R_ii 5′-CTATATTTGGGAAAGGCGCATGAATACC-3′. The TOXIN PCR cycling conditions were as follows: Initial denaturation at 94 °C for 30 sec; 30 cycles of 94 °C for 30 sec; 55 °C for 30 sec; 68 °C for 1.20 min; Final extension at 68 °C for 5 min; Hold: 4–10 °C. PCR products were stored at 4 °C and visualized on a 1.6% agarose gel stained with SYBR™ safe DNA gel stain (ThermoFisher Scientific). Cultures positive for the *toxA* gene were banked in brain–heart infusion broth (BHI) + 50% glycerol solutions and stored at –80 °C for further analysis.

### Antibiotic susceptibility testing

Antibiotic susceptibility was determined using the disk-diffusion method. Frozen stocks were thawed and inoculated onto CBA plates, followed by incubation at 30 °C for 72 hours. A loopful of colony material was inoculated into 3 mL (∼5 mm depth) of Todd Hewitt broth in a flat-bottomed vial, lined with approximately 1 mL of tryptic soy agar. Liquid cultures were statically incubated for 48 hours at 37 °C, thereafter cultures were resuspended and assessed visually for turbidity. A 1:1 dilution of the culture was made in Todd Hewitt broth and further statically incubated for 3–4 hours or until an increase in turbidity was apparent using McFarland standards; this allowed the *O. hominis* growth to recover. A volume of 100 µL of the diluted broth was inoculated onto CBA plates using the spread plate method. Three replicates were included for each strain. Antibiotic disks (Oxoid, ThermoFisher Scientific) of amoxicillin (25 U), ampicillin (10 U), vancomycin (5 U), polymyxin B (300 U), and penicillin G (1 U) were placed onto the agar surface using sterile forceps. Plates were incubated for 48 hours at 30 °C under microaerobic conditions. Growth was evaluated and zone sizes were measured using callipers.

Nitrocefin disks (Remel, ThermoFisher Scientific) were used according to the manufacturer’s instructions. In brief, disks were heavily inoculated with bacterial growth from solid media and incubated for 5 minutes in air at room temperature. A colour change from white to pink indicated a positive result for β-lactamase production, while no colour change indicated a negative result. Isolates were tested in triplicate.

### DNA extraction and sequencing

Confirmed *O. hominis* isolates were cultured on CBA and incubated at 30 °C for up to 120 hours for DNA extraction. Genomic DNA was extracted from pure bacterial cultures using the Wizard HMW DNA extraction kit (Promega, USA). The protocol was performed according to the manufacturer’s instructions with some modifications. Briefly, colonies were scraped from plates and pelleted in PBS by centrifugation at 16,000 × g for 2 minutes. The supernatant was discarded and the cell pellet was resuspended in PBS. Resuspensions were incubated at 85 °C for 5 minutes. 500 µL lysis buffer was added along with 6 µL RNase A and the mixture was incubated at 37 °C for 30 minutes. 20 µL of Proteinase K was added and the samples were incubated at 56 °C for 15 minutes. Protein was precipitated using protein precipitation solution. Thereafter, the samples were centrifuged at 16,000 × g for 10 minutes at room temperature. The supernatant was carefully transferred into room temperature isopropanol for DNA precipitation. The samples were centrifuged at 16,000 × g for 2 minutes to pellet the DNA. The DNA pellet was washed with 70% ethanol and the samples were centrifuged at 16,000 × g for 2 minutes before allowing the DNA pellet to air dry at room temperature. The DNA was left to resuspend in 50 µL of DNA resuspension buffer at room temperature. DNA was stored at 4 °C until ready for further analysis.

The sequencing library was prepared with the native barcoding kit 24 v12 (SQK-NBD112.24; Oxford Nanopore Technologies) according to the manufacturer’s guidelines. Briefly, DNA was FFPE repaired and end-prepped/dA-tailed, then a unique dT-tailed barcode adapter was ligated on the dA-tailed template. Barcoded samples were pooled and sequencing adapters were ligated. The library was sequenced on a Oxford Nanopore MinION platform using a flow cell R10.4 (FLO-MIN114; Oxford Nanopore Technologies).

### Genome analysis

Base calling and barcode removal was undertaken using the SUP model in Guppy v6.5.7 (GPU). Reads were trimmed using Prowler (17) to a Q20 average in a 1,000-nucleotide window, discarding reads of length <1 kb. Preliminary assemblies were generated with Flye v2.9.2 (18) using default settings. Genomes were circularised and rotated to the origin of replication as described previously (19). The genome was polished with Medaka v1.9.1 using trimmed reads. Finally, the genome was annotated with Bakta v1.8.1 (20). Due to poor performance of SA-OH-C4 specifically, an alternative pipeline was used in this case: AssembleBAC-ONT, developed by Andries van Tonder (https://github.com/avantonder/assembleBAC-ONT). Briefly, Artic guppyplex aggregated pre-demultiplexed reads from Guppy. Long reads were filtered using Filtlong, discarding reads of <5 kb. As with the other genomes, AssembleBAC-ONT assembled reads with Flye, polished with trimmed reads using Medaka, and annotated with Bakta.

Antimicrobial resistance genes were identified using NCBI’s AMRFinderPlus tool v3.12.8 (21) and the Comprehensive Antibiotic Resistance Database (CARD) Resistance Gene Identifier (RGI) tool v1.2.1 (22). Pangenome analysis was performed using Panaroo (23). Annotated assemblies of South African, Thai, and Australian *O. hominis* isolates were included in pangenome analysis.

Panaroo was used with default settings in the strictest mode and generated a core genome alignment (all genes present in 98% of isolates) using the ClustalW multiple sequence aligner. The core genome alignment tree (Figure 1a) was calculated in MEGA11 and visualised using ggtree in R v.4.4.2. Evolutionary history was inferred by using the Maximum Likelihood method and Tamura– Nei model. The bootstrap consensus tree inferred from ten replicates was taken to represent the evolutionary history of the taxa analysed. Branches corresponding to partitions reproduced in less than 50% bootstrap replicates are collapsed. The percentage of replicate trees in which the associated taxa clustered together in the bootstrap test are shown next to the branches. Functional prediction of core genes was performed using eggNOG-mapper (24) with bacteriophage detection using PHASTEST (25) and IS element prediction with ISEScan v1.7.2.3 (26). Comparison of amino acid sequences was undertaken with MMseqs2 v17 (27) using default settings with a custom database comprising all 99,432 RefSeq genomes (28) that belonged to a genus present in at least 1% abundance in at least one sample in the reported microbiome analysis, plus the top 20 genera reported for the full cohort (13). Further details are reported in the Supplementary Methods.

**Figure 1:**
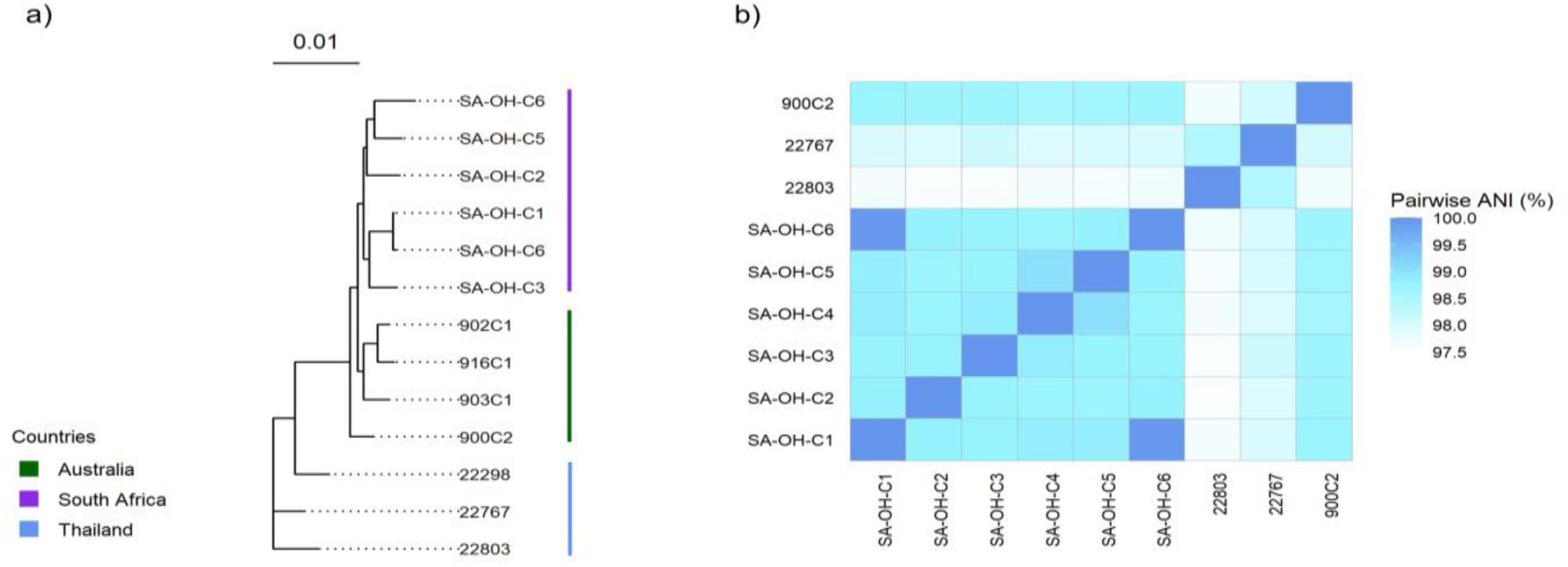
a) Maximum likelihood phylogenetic tree of core genes from South African (purple), Thai (blue), and Australian (green) *O. hominis* genomes; b) Heatmap of pairwise average nucleotide identity.

### Microbiome analysis

Using the 16S rRNA dataset described above, samples were selected from 23 infants with the highest predicted *O. hominis* abundance (*O. hominis* carrier group, timepoint 2). The closest preceding sample that had no *O. hominis* reads was selected as a pre-colonisation comparator (*O. hominis* carrier group, timepoint 1). Non-carriers of *O. hominis* were identified in the dataset based on an absence of *O. hominis* reads from birth to 30 months of age, and 23 of these infants were randomly assigned as paired controls. Age matched samples were selected (non-carrier group, timepoint 1 and 2). One infant had *O. hominis* in all their preceding samples, so timepoint 1 was omitted for this and the paired control. The negative controls associated with these sequencing runs were included in the analysis for decontamination purposes. The selection criteria are further illustrated in Supplementary Figure S2, and metadata of the carrier and non-carrier groups in Supplementary Table S1.

Data from these 90 samples and 14 negative controls were cleaned using Mothur v1.44.3 (29). In brief, contiguated read pairs were removed if they had any ambiguous bases, homopolymers of length >8, or a total length >260bp, and following alignment they were trimmed to the expected V4 region of the 16S rRNA gene. Chimeras were identified using the chimera.vsearch function with dereplication and removed from downstream analysis (in total 7.9% of reads were removed). Reads were classified against the SILVA database v132 (30) and those identified as Chloroplast, Mitochondria, Archaea, Eukaryota, or of unknown Kingdom were removed. Finally, actual sequence variants (ASVs) comprising only one read were discarded. The ASV matrix from Mothur was converted to a phyloseq object in R v4.2.1 (31) in order to run Decontam v1.12 (32) using the prevalence method with a threshold of 0.5. Further details of decontamination are described in the Supplementary Methods.

Alpha diversity metrics (coverage, observed ASVs, Inverse Simpson index) were calculated in Mothur with rarefaction to the lowest sample depth, 10,084. Paired t-tests in R were used to compare ASV richness between groups. Due to non-normal distribution, diversity scores were compared with the Mann–Whitney U test.

ASV network analysis was undertaken for well represented taxa, i.e. all 60 ASVs with a relative abundance of >1% in at least one sample and excluding singletons. Network construction and visualization used NetCoMi v1.2.0 (33) with the SparCC association measure for a fully connected (dense) network, identifying hub nodes by the eigenvector method. Edges were filtered by weight <0.25. ASVs were classified using the RefSeq Targeted Loci database.

## Results

### Identification and recovery of *O. hominis* from nasopharyngeal samples

*O. hominis* reads were detected in 629/6025 nasopharyngeal samples from 204/605 children in the microbiome dataset (10.4% NP samples or 33.7% of participants) as well as in 14/330 sputum samples from 12/239 infants (4.2% of sputum samples or 5% of participants). In six of these sputum samples (from five infants), *O. hominis* was also detected in the nasopharynx on the same date or immediately preceding it.

Nasopharyngeal samples were selected for isolation attempts based on the following criteria: (i) longitudinal carriage of *O. hominis* (presence of *O. hominis* reads in more than one nasopharyngeal sample from the same child), (ii) availability of child age data, and (iii) expected *O. hominis* abundance greater than 1%. 34 NP samples fulfilled these criteria and were targeted for isolation attempts; however, *O. hominis* was successfully isolated from only four of these samples. Two additional isolates (SA-OH-C1 and SA-OH-C5) were cultivated from samples that were identified using data from a prior metagenomic study (PRJNA727021, manuscript in preparation).

In summary, six *O. hominis* isolates were recovered and sequenced from samples collected between 2013 and 2016. The age of the children at the time of sampling ranged from 238 days (7.8 months) to 761 days (>2 years of age), and the relative abundance of *O. hominis* in the NPS was between 1–6%. All isolates were obtained from swabs collected during routine study sampling, except for two infants, C1 and C5, who were sampled during acute respiratory illness (Table 1).

**Table 1:**
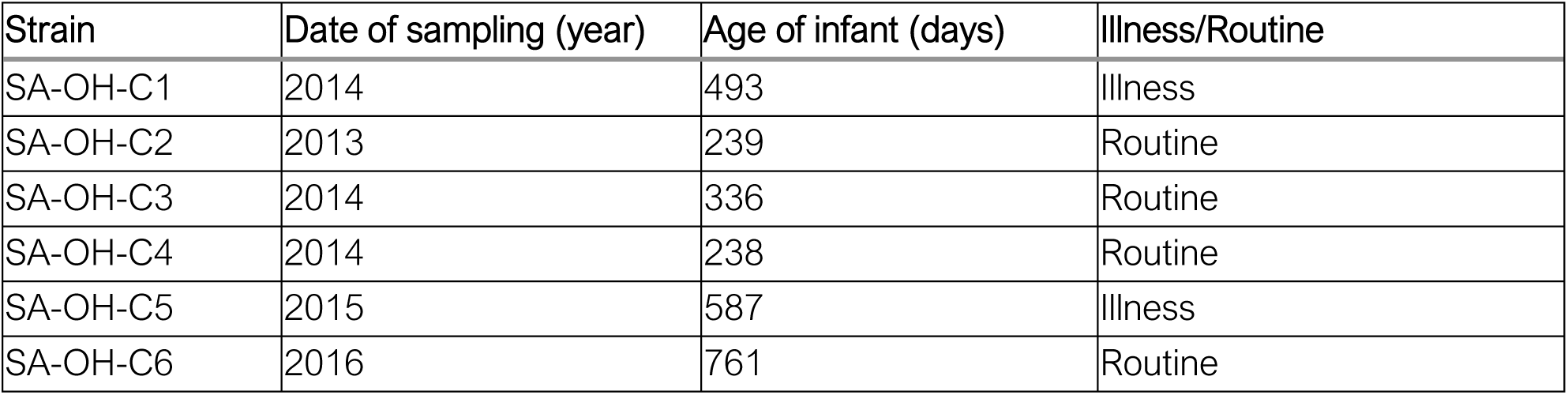
Sample information for *O. hominis* isolates.

Colonies typical of *O. hominis* morphology as described by Lawrence et al (2019) were observed; colonies appeared greyish, glistening, convex, and circular with smooth edges (12). Isolate SA-OH-C6 exhibited a consistently mucoid colony morphology. A subculture of a single colony could yield a variety of colony sizes, ranging from punctiform (pinprick-sized, less than 1 mm in diameter) to larger colonies up to 3 mm in size. Uniformly sized colonies of ∼1 mm in diameter were also apparent. Notably, punctiform colonies from mixed morphology plates typically become apparent after prolonged incubation and increased in frequency with extended incubation periods. *O. hominis* colonies also exhibited α-haemolysis on CBA plates after extended incubation (>96 hours). This haemolysis was most pronounced under the primary streak. Haemolytic activity was not observed for the smaller, punctiform colonies. Growth and appearance of colonies became more luxuriant following passage in shallow Todd Hewitt broth.

### Characteristics of South African *O. hominis* genomes

As summarized in Table 2, all genomes were approximately 2 Mb in size with sequencing coverage ranging from 17× to 55×. Average GC content was 35.7% and there were approximately 2,000 genes in each genome; SA-OH-C2 appears to have more genes than the other genomes, an artificial inflation caused by sequencing artefacts, however this did not affect downstream analysis due to the successful recovery of fragmented genes by Panaroo. Single, circular chromosomes were generated for all genomes except for SA-OH-C4 which assembled into 6 contigs, one of which was identified as a putative plasmid and another as a 33 kb bacteriophage.

**Table 2:** *O. hominis* genome characteristics.

| Genome | Accession number | Total length (Mbp) | No. of contigs | Complete | Mean coverage (x) | GC content (%) | Total genes (CDS) | Plasmids |
| --- | --- | --- | --- | --- | --- | --- | --- | --- |
| SA-OH-C1 | ERS21330455 | 2.05 | 1 | Yes | 55 | 35.75 | 1991 (1934) | 0 |
| SA-OH-C2 | ERS21330459 | 2.05 | 1 | Yes | 23 | 35.69 | 2183 (2123) | 0 |
| SA-OH-C3 | ERS21330456 | 2.01 | 1 | Yes | 51 | 35.69 | 1900 (1840) | 0 |
| SA-OH-C4 | ERS21330457 | 1.99 | 6 | Contigs | 50 | 35.71 | 1965 (1908) | 1 |
| SA-OH-C5 | ERS21330460 | 2.05 | 1 | Yes | 17 | 35.49 | 2045 (1987) | 0 |
| SA-OH-C6 | ERS21330458 | 2.05 | 1 | Yes | 46 | 35.73 | 2089<br>(2030) | 0 |

The core genome tree (Figure 1a) illustrates the relationship between the genomes from South African and Australian isolates as well as genomes from the Mae La cohort in Thailand. The Australian and South African genomes appear to be more closely related, while the genomes from Thailand are more distantly related. Pairwise average nucleotide identities (ANI) were >97% between the genomes (Figure 1b), confirming that they are members of the same species.

The functions of the accessory and core genes of *O. hominis* were investigated using EggNOG-mapper with supplementary assignment of bacteriophage and IS elements using PHASTEST and ISEScan. Clusters of orthologous groups (COGs) were assigned genes where possible, and the distribution of gene counts across the COG categories is presented in Figure 2. Approximately 800 COGs were identified for the accessory genome and 900 COGs for the core genome, although a large proportion of the accessory genes (215/800, 26.87%) and core genes (179/900, 19.89%) could not be functionally annotated. The accessory genome annotations were dominated by groups associated with replication and repair, cell wall biogenesis, and bacteriophage. The first and second categories are somewhat inflated as “replication and repair” includes genes from mobile elements such as integrase/recombinases and nucleases and “cell wall biogenesis” includes 26 orphan tips (pseudogenes) associated with a large RHS protein. In contrast, genes linked to secondary metabolite biosynthesis and catabolism, signal transduction, nucleotide metabolism, and cell motility were poorly represented in the accessory genome.

**Figure 2:**
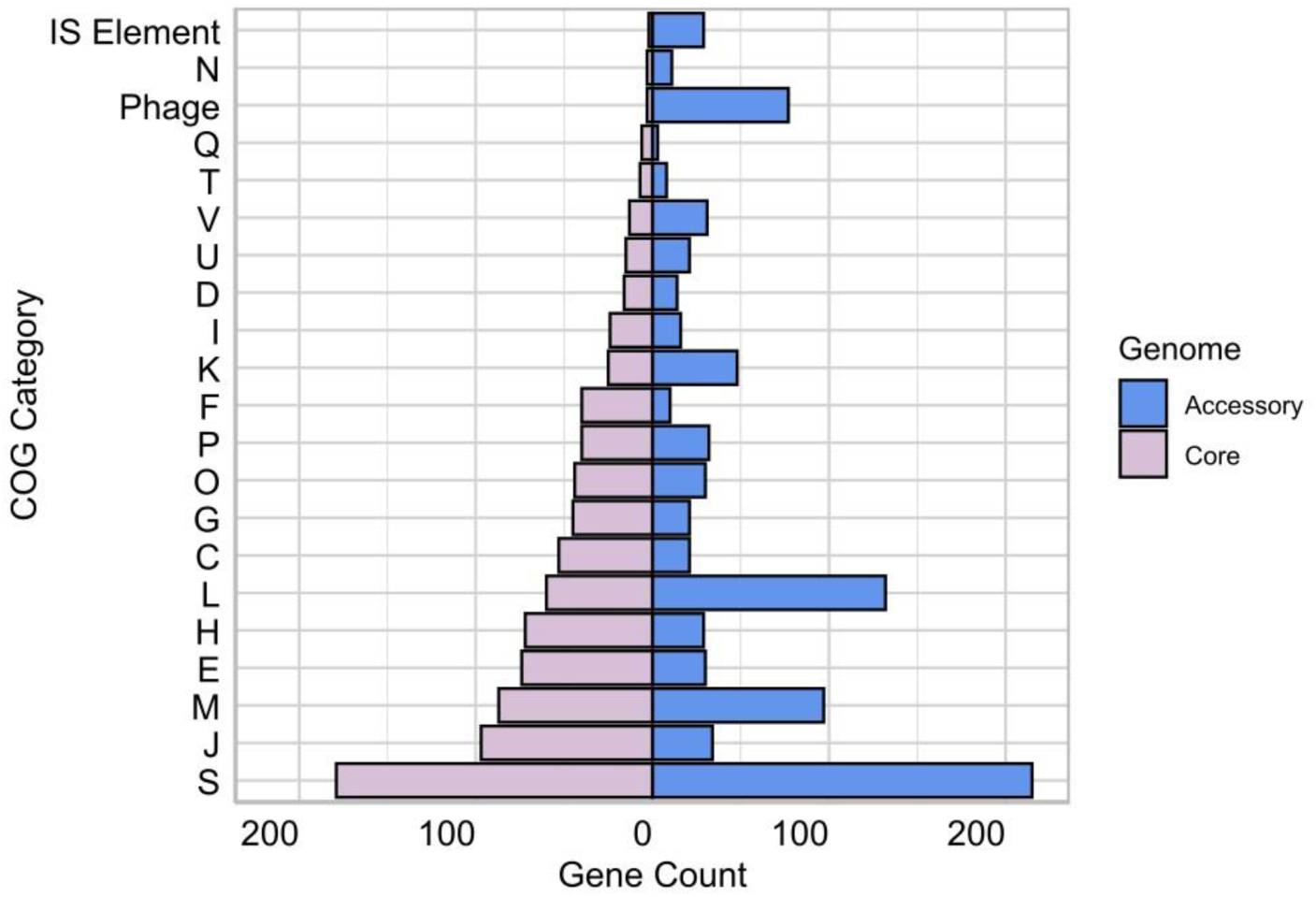
Summary of accessory genome COG groups. C: Energy production and conversion; D: Cell cycle control, cell division, chromosome partitioning; E: Amino acid transport and metabolism; F: Nucleotide transport and metabolism; G: Carbohydrate transport and metabolism; H: Coenzyme transport and metabolism; I: Lipid transport and metabolism; J: Translation, ribosomal structure, and biogenesis; K: Transcription, L: Replication, recombination, and repair; M: Cell wall/membrane/envelope biogenesis; N: Cell motility, O: Post-translational modification, protein turnover, chaperone functions; P: Inorganic ion transport and metabolism; Q: Secondary metabolites biosynthesis, transport, and catabolism; T: Signal transduction; U: Intracellular trafficking, secretion, and vesicular transport; S: Function unknown; V: Defence mechanisms; and X: Mobilome.

In the core genome, most genes were associated with translation and ribosome function, cell membrane biogenesis, and metabolism of amino acids and coenzymes. Like the accessory genome, core genes related to cell motility and signal transduction were the least represented among the categorized genes in *O. hominis*. No genes were assigned to Z (cytoskeleton), Y (nuclear structure), X (mobilome), W (extracellular structures), R (only general function prediction), and A (RNA processing and modification) COG categories.

All six *O. hominis* genomes include novel prophages. SA-OH-C2 encodes three prophage regions, while SA-OH-C1, -C3, -C4, and -C6 have two, and SA-OH-C5 has one. Where multiple phages are present, they are dissimilar to one another, except for SA-OH-C4 which includes one intact bacteriophage chromosome and at least one truncated copy of the same prophage inserted into the bacterial chromosome. The variable presence of this prophage may be responsible for the assembled genome’s fragmentation into contigs.

### Antimicrobial resistance potential in *O. hominis*

The AMR Finder and CARD resistance gene identifier tools were used to screen for AMR genes in the genomes and isolates were evaluated for β-lactamase production and susceptibility to polymyxin B, vancomycin, amoxicillin, ampicillin, and penicillin G (Table 3).

**Table 3:**
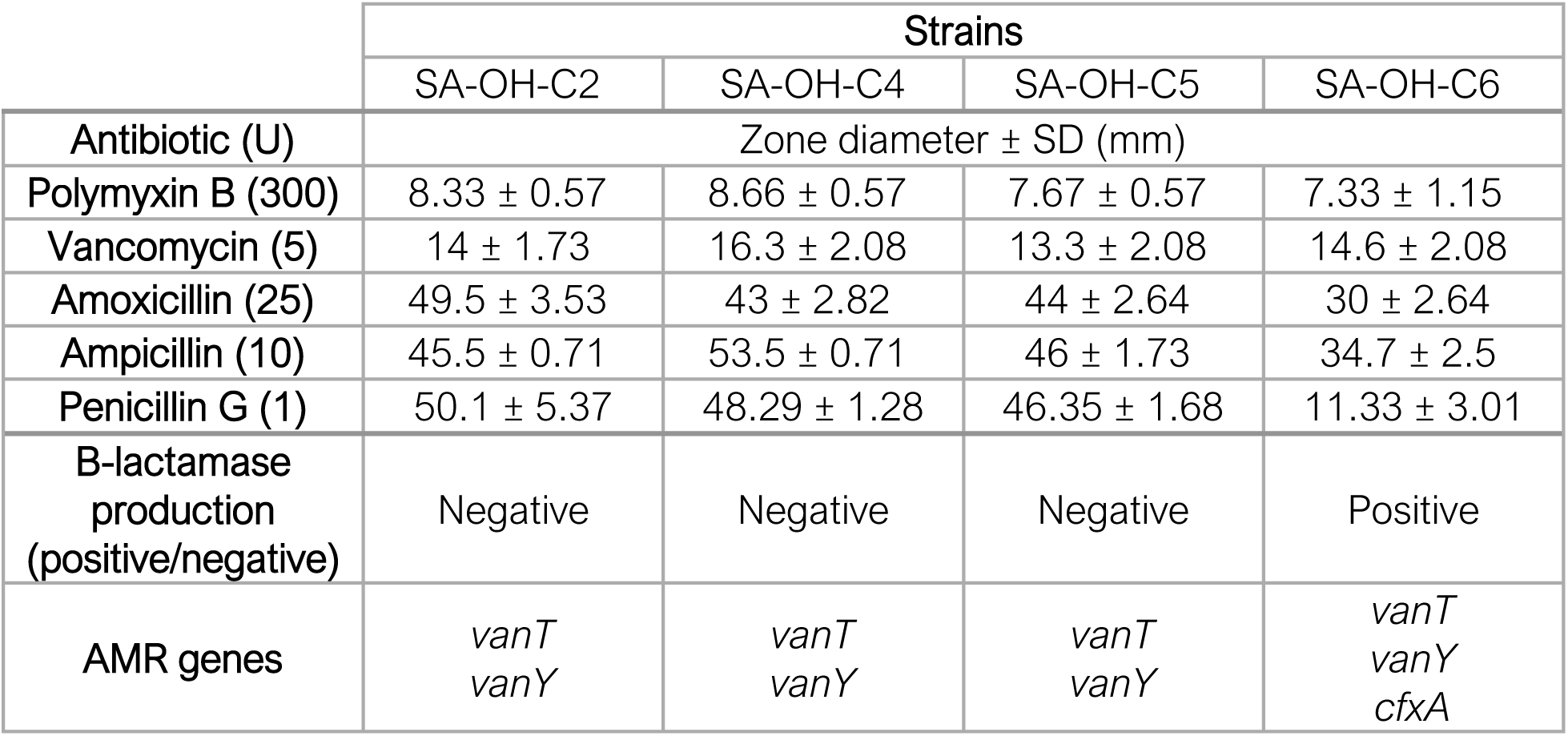
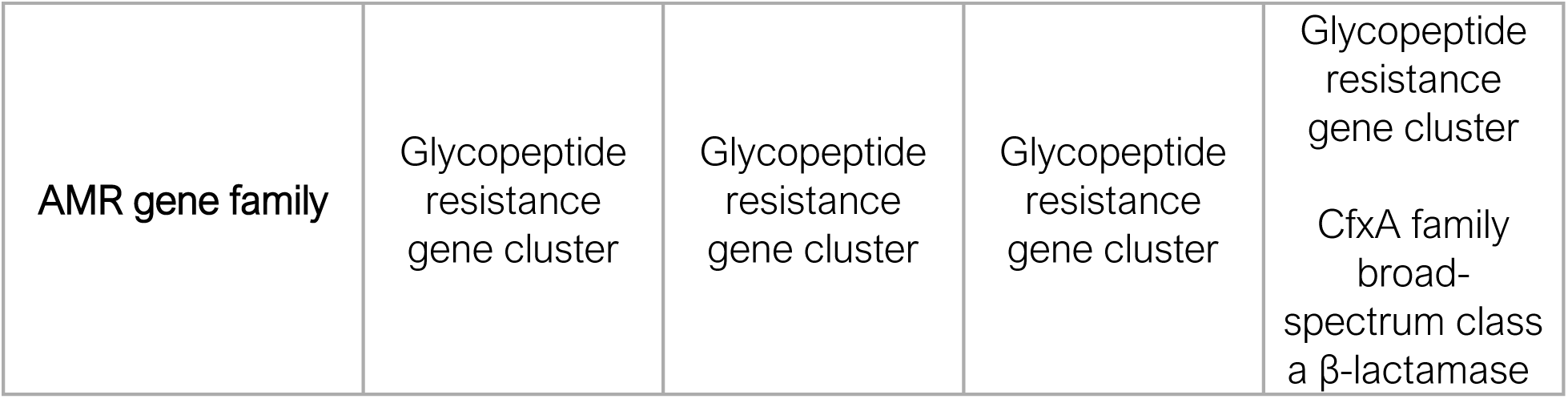
Antibiotic resistance potential screening of South African *O. hominis* isolates.

|  | Strains |  |  |  |
| --- | --- | --- | --- | --- |
|  | SA-OH-C2 | SA-OH-C4 | SA-OH-C5 | SA-OH-C6 |
| Antibiotic (U) | Zone diameter ± SD (mm) | | | |
| Polymyxin B (300) | $8.33 \pm 0.57$ | $8.66 \pm 0.57$ | $7.67 \pm 0.57$ | $7.33 \pm 1.15$ |
| Vancomycin (5) | $14 \pm 1.73$ | $16.3 \pm 2.08$ | $13.3 \pm 2.08$ | $14.6 \pm 2.08$ |
| Amoxicillin (25) | $49.5 \pm 3.53$ | $43 \pm 2.82$ | $44 \pm 2.64$ | $30 \pm 2.64$ |
| Ampicillin (10) | $45.5 \pm 0.71$ | $53.5 \pm 0.71$ | $46 \pm 1.73$ | $34.7 \pm 2.5$ |
| Penicillin G (1) | $50.1 \pm 5.37$ | $48.29 \pm 1.28$ | $46.35 \pm 1.68$ | $11.33 \pm 3.01$ |
| β-lactamase production (positive/negative) | Negative | Negative | Negative | Positive |
| AMR genes | *vanT*<br>*vanY* | *vanT*<br>*vanY* | *vanT*<br>*vanY* | *vanT*<br>*vanY*<br>*cfxA* |
| AMR gene family | Glycopeptide resistance gene cluster | Glycopeptide resistance gene cluster | Glycopeptide resistance gene cluster | Glycopeptide resistance gene cluster<br>CfxA family broad-spectrum class a β-lactamase |

An incomplete set of genes associated with vancomycin resistance (v*anT* and *vanY)* were identified in all South African strains. Two strains, SA-OH-C1 and SA-OH-C6, carry the c*fxA* gene, which encodes a class A2 β-lactamase that may confer broad activity against β-lactam antibiotics. Strains SA-OH-C1 and SA-OH-C3 were not subjected to antibiotic susceptibility testing due to issues with sample viability and contamination; all other isolates were tested in triplicate. Zone diameters are reported in Table 3. Vancomycin inhibition zones were similar across the isolates, ranging from 13.3 mm to 16 mm. Although breakpoints are not established for amoxicillin and ampicillin in this species, all isolates exhibited large inhibition zones for these β-lactam antibiotics. Notably, SA-OH-C6, which carries the c*fxA* gene, exhibited a reduced inhibition zone for penicillin G compared to the other isolates (11.33 ± 3.01 mm compared to a mean of 48.25 mm). This strain also yielded a 13–20 mm and an 11–18 mm reduction in zone diameter for ampicillin and amoxicillin antibiotics, respectively, and was the only isolate with a positive β-lactamase production test.

### Exchange of virulence determinants: LPS

The structure and composition of the lipopolysaccharide (LPS) biosynthesis loci in the South African genomes were analysed and compared to available LPS clusters from public *O. hominis* data. The alignment (Figure 3) of the LPS locus illustrates its composition and organization. As described previously (12), the initial 14 kb and two terminal sugar transferase genes are highly conserved across the LPS types, offering a conserved site for variation through homologous recombination.

**Figure 3:**
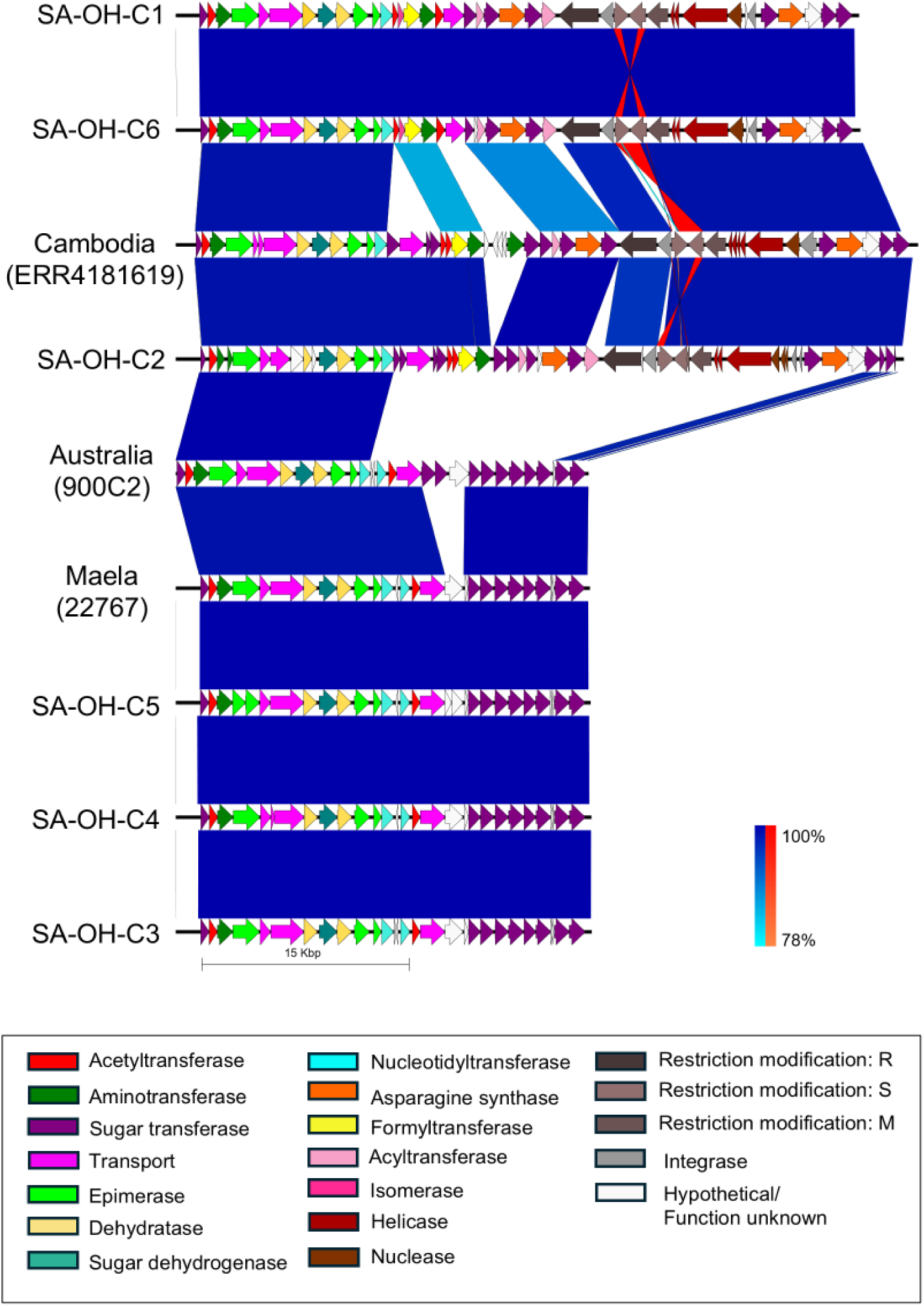
Alignment of lipopolysaccharide biosynthesis clusters in the South African genomes and *O. hominis* from Cambodia, Australia, and Thailand.

SA-OH-C3, -C4 and -C5 possess a LPS cluster identical to that of the Thai genome OH-22767 (accession no. GCF_900538225.1), demonstrating that the LPS is decoupled from the structure of the core genome tree (Figure 1a). The LPS clusters from SA-OH-C1, -C2, and -C6, although similar to the Cambodian LPS type (derived from accession no. ERR4181619), exhibit differences in the central region of the cluster. SA-OH-C1 and -C6 include a novel transport gene, possibly a novel flippase, as well as distinct glycosyltransferases and a unique acyltransferase. Compared to SA-OH-C1 and -C6, SA-OH-C2 is more similar to the Cambodian LPS type with shared transport genes, however, -C2 has an additional acyltransferase and lacks the aminotransferase.

In South African isolates SA-OH-C1, -C2, and -C6 the LPS gene cluster also encompasses a 15 kb putative non-autonomous transposon. Flanked by two tyrosine type integrase/recombinase genes, the transposon contains a YhcG nuclease gene, two helicase genes, and type I restriction enzyme subunits SMR. Although the complete element has not been observed outside of *O. hominis*, the individual genes have a high amino acid identity (>80%) to others within the family, primarily from *Chryseobacterium spp*. The exception is the YhcG nuclease and one helicase, which share 99% and 96% amino acid identity, respectively, with genes from a mobile element in *Sphingobacterium mizutaii* strain SM2 (acc. no. NZ_JACLGQ010000002), a clinical isolate from China. The species *S. mizutaii* was originally classified as *Flavobacterium*.

### The nasopharyngeal microbiome of *O. hominis* carriers differs from non-carriers

Using the screened 16S rRNA gene data as a starting point, matched samples from 23 *O. hominis* carriers with >1% relative abundance (sampled pre- and post-colonisation) and 23 non-carriers were examined. To overcome the confounding influence of age on microbiome composition, samples from *O. hominis* carriers and non-carriers were well matched, with a median difference in age of 2 days (maximum difference 37 days) (Supplementary Figure S3). Alpha diversity measures show increased diversity (Inverse Simpson index) between timepoint 1 and 2 for the *O. hominis* carrier group (p=0.005) with no significant change in the non-carrier group. Furthermore, the diversity scores of samples with *O. hominis* are higher than age matched samples from non-carriers (p=0.032) (Figure 4). The median increase in community richness (observed ASVs) over time for *O. hominis* carriers was 14.1 ASVs, but the increase was not significant.

**Figure 4:**
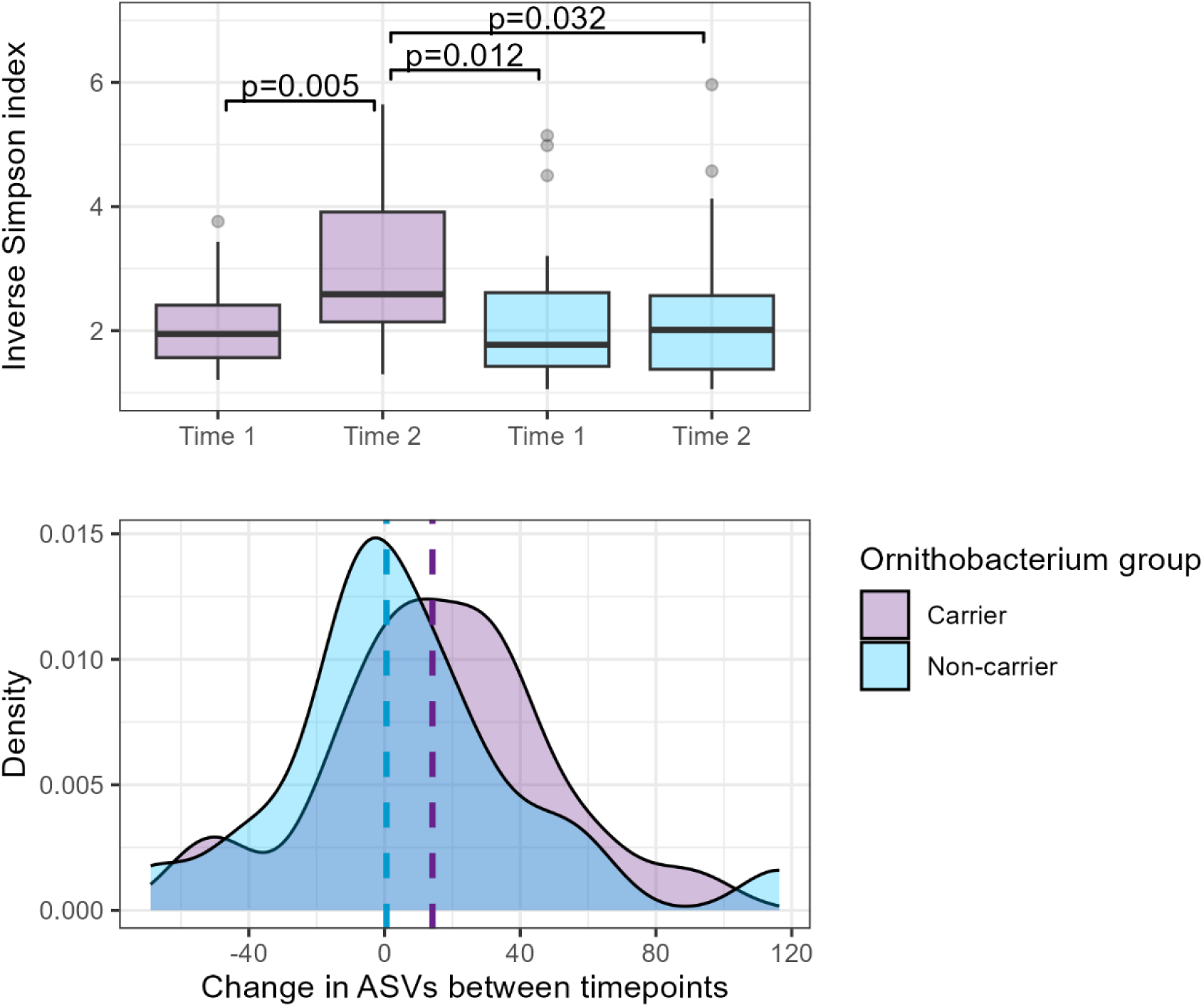
Boxplot (top) of Inverse Simpson diversity scores for *O. hominis* carriers (purple) and non-carriers (blue). The *O. hominis* carrier group at timepoint 2 is significantly more diverse than at the earlier timepoint and in comparison to non-carriers. Distribution plot (bottom) of the change in community richness per infant between timepoints, with the group median indicated by dashed line (*O. hominis* carriers +14.1 ASVs, non-carriers +0.6 ASVs).

A correlation network was calculated using 60 key ASVs, encompassing 96.4% of total reads. As illustrated in Figure 5a, the ASVs were categorized into two clusters: Cluster 1 (in purple) that includes a well-connected subcluster with two hub nodes, *O. hominis* and *Helcococcus* sp, and Cluster 2 (in blue) with one hub node, an unclassified Lachnospiraceae.

**Figure 5:**
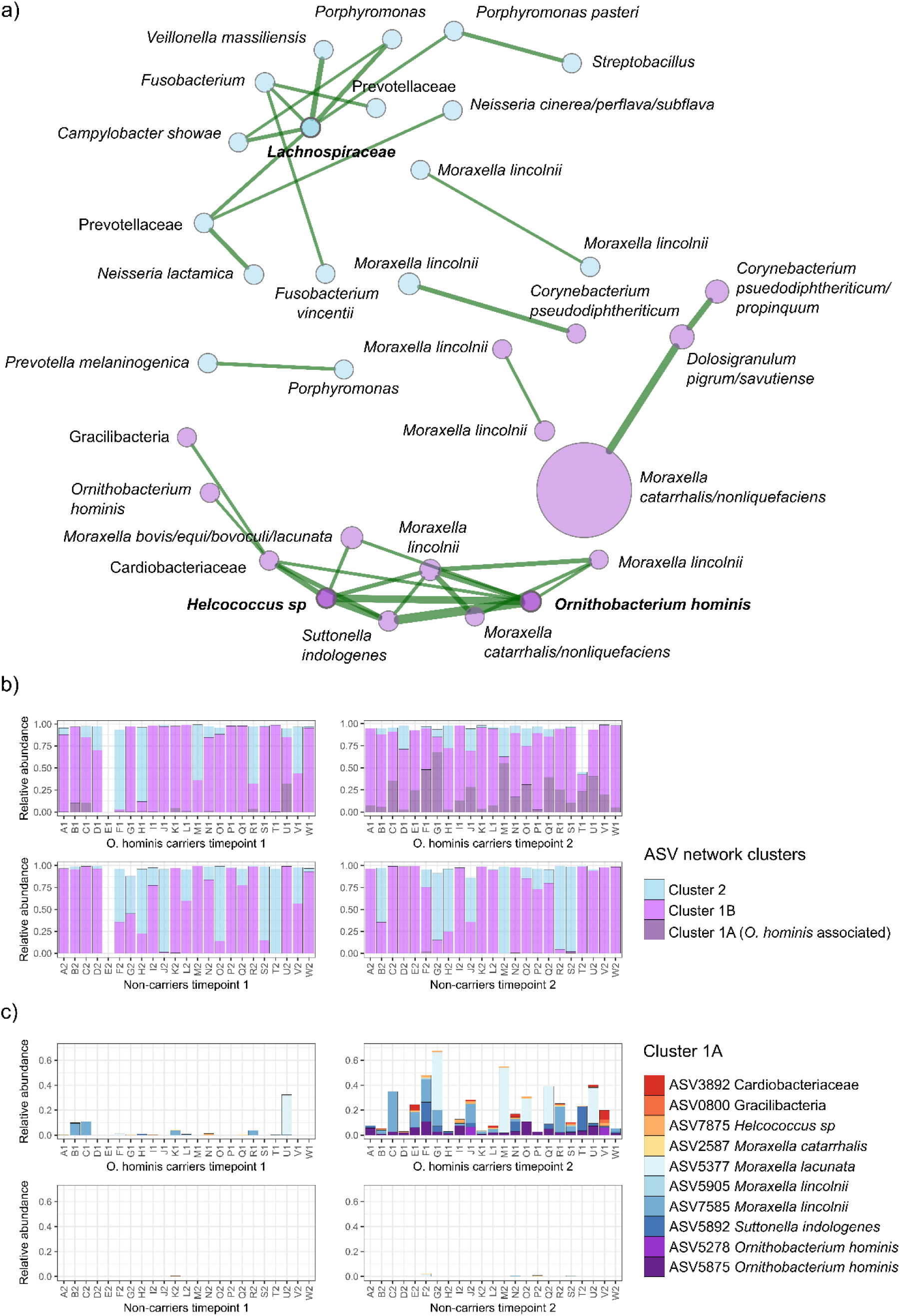
a) SparCC network diagram illustrating connected ASVs within cluster 1 (purple nodes) and cluster 2 (blue nodes), and all positive associations with weight >0.25 (green edges). Three hub nodes were identified (in bold): *O. hominis, Helcococcus* sp, and an unclassified Lachnospiraceae. Node sizes are scaled by normalized counts. b) Relative abundance of the ASV clusters at two timepoints in *O. hominis* carriers (A1-W1) and non-carriers (A2-W2), with Cluster 1 divided into the *O. hominis-*associated subcluster 1A and non-associated ASVs (1B). c) Composition of subcluster 1A.

Subcluster 1A comprises ASVs identified as *O. hominis*, an unclassified Cardiobacteriaceae, unclassified Gracilibacteria, *Helcococcus* sp*, M. catarrhalis/nonliquefaciens, M. lacunata/bovis/equi/bovoculi, M. lincolnii,* and *S. indologenes*. The remainder of Cluster 1 comprises ASVs that are not significantly correlated with *Ornithobacterium*, including *Corynebacterium propinquum/pseudodiphtheriticum, Dolosigranulum pigrum/savutiense*, and a highly abundant *M. catarrhalis/noniquefaciens* ASV.

Cluster 2 includes several oral-associated taxa including *Fusobacterium*, *Porphyromonas* and *Veillonella* that may reflect saliva intrusion into the nasopharyngeal sampling site.

Visualising the clusters present in the *O. hominis* carriers and non-carriers at the pre- and post-colonisation timepoints (Figure 5b, 5c) illustrates that some *Ornithobacterium*-associated taxa are present before the acquisition of *O. hominis* and that they are less common in the non-carrier group overall. The most abundant of these are the *M. lincolnii*-like ASVs (Figure 5c).

Isolates SA-OH-C2, SA-OH-C3, and SA-OH-C6 were recovered from infants O1, Q1, and J1, respectively (Figure 5b, c). Isolates SA-OH-C1, -C4 and -C5 were obtained from infants not included in the paired carrier/non-carrier analysis. During attempts to isolate *O. hominis* from primary samples, bacterial co-colonisers were also recovered at 30 °C under microaerobic conditions. One isolate of *Helcococcus,* SA-HE-C1, was sequenced, yielding a single circular chromosome of 1.4 Mbp and 28.36% GC content. SA-HE-C1 shares <74% average nucleotide identity to members of the five published species and <95% 16S rRNA gene identity (Table 4), and its 16S rRNA gene sequence matches that of the *Helcococcus* sp ASV above.

**Table 4:** Average nucleotide identity and 16S rRNA gene identity to the five known species of *Helcococcus*, as defined by GTDB. The two unnamed species, from metagenomes, lack an assembled 16S rRNA gene.

|  | Pairwise ANI<br>to SA-HE-C1 | 16S rRNA gene identity to<br>SA-HE-C1 |
| --- | --- | --- |
| *Helcococcus kunzii* ATCC-51366,<br>GCF000245755 | 73.5%<br>(across 422 kbp) | 94.1% |
| *Helcococcus ovis* KG40,<br>GCA004524815 | 73.6%<br>(across 426 kbp) | 93.9% |
| *Helcococcus sueciensis* DSM-17243,<br>GCF000423145 | 73.5%<br>(across 424 kbp) | 94.3% |
| *Helcococcus* sp023418535,<br>GCA023418535 | 72.7%<br>(across 368 kbp) | Gene not present |
| Helcococcus sp023442495,<br>GCA023442495 | 71.7%<br>(across 303 kbp) | Gene not present |

### *O. hominis* as a reservoir of adaptive genes from the microbial community

The *O. hominis* genomes were compared to a custom database of 99,432 reference genomes including every available species from a genus with sample abundance >1% in the microbiome analysis, and the 20 most abundant genera in the full cohort, to identify genes that may be shared with other nasopharyngeal species. Five blocks of genes were identified altogether, with individual gene similarity >99% to other genera.

Evidence of inter-species sharing of antimicrobial resistance genes is provided by the well-studied transposon Tn4555, a 10 kb transposon carrying the *cfxA* β-lactamase gene, found in SA-OH-C1 and -C6. Although this transposon is widespread, the nucleotide sequence present in *O. hominis* is >99.9% identical to that observed in *Prevotella scopos, P. denticola, P. melaninogenica*, and *P. distasonis* genomes, all of which are oral species of *Prevotella*.

Adhesins are another way in which bacteria can adapt to the host environment. All *O. hominis* genomes to date include a variable set of at least 20 *Fibrobacter succinogenes* domain (FSD) genes, many of which also contain fibronectin-binding domains and have an associated putative phase variation mechanism. Large complements of FSD genes are known in only a few other bacteria, such as *Fibrobacter succinogenes* (a rumen commensal) from which they are named, *Chryseobacterium nematophagum* (a pathogen of nematodes), and *Elizabethkingia anophelis* (a mosquito gut commensal). A block of three exogenous genes in SA-OH-C1 and -C6 consist of a protease, a putative preprotein, and an FSD gene, altogether sharing >99.9% nucleotide identity with *Bacteroides heparinolyticus* strain F0111. This implies that adhesins such as FSD proteins may be shared between species.

Finally, RHS toxin genes encode a large protein apparatus enclosing a C terminal toxin that is delivered to a prokaryotic or eukaryotic target cell by a type VI secretion system (34). The toxin tips are variable and change through a displacement mechanism, generating an array of pseudogene tips and immunity genes (35). All six South African *O. hominis* isolates carried a RHS toxin tip and immunity gene within their array which has also been observed in a group of *Bergeyella cardium* isolates (36) but has not been found in *O. hominis* from elsewhere. The *B. cardium* references possess a similarly structured RHS array, although the primary RHS protein is of a different sequence to that of *O. hominis,* indicating that toxin tips can also be shared between species.

## Discussion

*O. hominis* was successfully isolated from six nasopharyngeal samples creating, to the best of our knowledge, the first cultured isolate collection of the species from African samples. Colony morphology and growth characteristics were consistent with published reports from Australia; haemolysis was observed, a delayed phenomenon also described in some strains of *O. rhinotracheale* (37); β-lactamase production and sensitivity to β-lactam antibiotics differed between strains. CfxA is a well characterised class A2 extended spectrum β-lactamase that hydrolyses penicillins and broad-spectrum cephalosporins. It is commonly present among oral bacteria of the class Bacteroidia, including *Prevotella*, *Porphyromonas*, and *Bacteroides* (38,39). In isolates SA-OH-C1, SA-OH-C6, and the previously published *O. hominis* genome OH-22803 from Thailand (11), the *cfxA* gene is present within transposon Tn4555 (40). The transposon insertion sites differ between the South African and Thai examples but are in a similar region of the chromosome: within 200kb of the terminus. The presence of *cfxA* appears to confer reduced susceptibility to β-lactam antibiotics, with the most profound effect observed against penicillin G.

By comparing infants who acquired *O. hominis* during the study period with those who were not colonised, we are able to identify differences in diversity and community membership that are associated with colonisation by this species. Community richness increases following colonisation with *O. hominis*, with a median increase of 14.1 ASVs. This is not simply a matter of increasing age, as it is not observed in matched non-carriers. Diversity also increases, with the *Ornithobacterium* colonised individuals having significantly more diverse communities than non-carriers of the same age. The increased inverse Simpson score is probably due to the establishment of not just one species, but several co-colonisers highlighted in the network analysis: *Suttonella, Helcococcus, Moraxella* spp, and an unclassified Gracilibacteria.

*Moraxella* is a dominant member of the nasopharyngeal microbiome in both groups. The genus appears to be slightly more abundant in *Ornithobacterium* carriers: mean 60.3% (timepoint 1) and 71.9% (timepoint 2), compared to 52.8% and 45.2%, respectively, for non-carriers. However, it is not homogenous, with *M. catarrhalis*-like ASVs present throughout the dataset but *M. lincolnii* and *M. lacunata*-like ASVs more markedly abundant in *Ornithobacterium* carriers. Based on the appearance of *M. lincolnii* before colonization with *O. hominis*, and the relative rarity of these ASVs among non-carriers, carriage of this species may be a prerequisite for *O. hominis* acquisition. The consistent co-occurrence of *Suttonella, Helcococcus*, and *Moraxella* with *O. hominis* could indicate a biofilm community that aids *Ornithobacterium* survival in the nasopharynx.

Correlation Cluster 1B, consisting of ASVs that are not part of the *Ornithobacterium* subcluster, includes protective or health-associated species of *Dolosigranulum* and *Corynebacterium* (8) known to inhibit pneumococcal/staphylococcal growth. Notably, *O. hominis* is inversely correlated with corynebacteria in this dataset, particularly *C. accolens*.

*O. hominis* is naturally competent and thus expected to acquire new genetic material through uptake from its environment. As a persistent coloniser of the upper respiratory tract, it may act as a reservoir of antibiotic resistance genes or virulence factors. Functional analysis of the genomes supports this, for example in the over representation of “replication and repair” genes originating from mobile elements. We assessed the potential contribution of genes from other species and identified several regions that may have been acquired from other species through mobile elements, such as the antimicrobial resistance associated Tn4555, and potentially via recombination of adhesion factors and neighbouring genes, such as those found in *B. heparinolyticus* or a RHS toxin tip from *B. cardium*. Interestingly, all matches were to oral species of bacteria rather than nasal colonisers. Recombination between nasal and oral bacteria has been described before, for example driving the mosaic penicillin-binding protein diversity of *S. pneumoniae* and thus its acquisition of β-lactam resistance (41).

A key virulence factor of Gram-negative bacteria is LPS which triggers an inflammatory response from the host immune system. As LPS is a highly immunogenic surface molecule, its biosynthetic gene clusters are common sites of recombination in bacteria. The South African genomes are more similar to one another than to those from elsewhere (Figure 1), but the close relationship between LPS biosynthetic gene clusters observed in different countries infers that these have been exchanged through recombination. The LPS operon appears to have been disrupted by an insertion event in three South African genomes, similar to a Cambodian example (Figure 3). The putative transposon is at the same location in different LPS gene clusters, yielding the same 135 bp insertion site duplication. This may be due to independent events at a conserved target site, or an historic insertion that was passed on through recombination at the LPS locus. There is no evidence for dynamic or variable presence of this mobile element in the raw long read sequence data from plate sweeps of multiple colonies, so it may no longer be functional. It is not known what the impact of an insertion on LPS biosynthesis may be.

Finally, isolate SA-HE-C1 of a novel *Helcococcus* species was recovered. The 1.4 Mbp genome shares <75% ANI and <95% 16S identity to all published species of the genus, thus we propose that it belongs to a new species, *Helcococcus ekapensis* (L adj, in reference to the origin of the isolate, Ekapa, the Xhosa name for Cape Town).

This study describes the genomes of the first *O. hominis* strains isolated in South Africa, and the differential nasopharyngeal microbiome profiles of infants who carry *O. hominis* compared to non-carriers. The main limitation of this study is the small number of isolates and genomes, which limits the ability to infer population structure. More isolates and genomes are required to address this limitation. Another limitation is the large fraction of hypothetical genes (on average 12% of genes) for which we cannot infer a function. This is perhaps a reflection of poor characterization of novel genes across the Weeksellaceae family (for example *Weeksella virosa* with 16% hypothetical genes, *Chryseobacterium indologenes* with 20%, or *O. rhinotracheale* with 25%) compared to well-studied species such as *Escherichia coli* for which fewer than 5% genes are uncharacterised.

## Ethical approval and consent to participate

Both this study (585/2015) and parent study (401/2009) received ethical approval from the Faculty of Health Sciences, Human Research Ethics Committee (HREC) of the University of Cape Town, South Africa. The relevant guidelines and regulations were followed during the performance of all experiments. Mothers participating in the parent study provided informed, written consent for enrolment of their infants at the time of delivery and annually.

## Author contributions

CCDA: formal analysis, investigation, data curation, writing - original draft preparation.

SJS: conceptualisation, funding acquisition, formal analysis, investigation, methodology, visualization, data curation, writing - original draft preparation.

SEB: investigation, supervision, writing - review and editing. SCW: resources, investigation, writing - review and editing. KSM: formal analysis, writing - review and editing.

HJZ: resources, writing - review and editing. MPN: resources, writing - review and editing.

FSD: conceptualization, funding acquisition, project administration, resources, supervision, writing - review and editing.

JP: conceptualization, funding acquisition, project administration, resources, supervision, writing - review and editing.

