## Supplementary methods for "Characterisation of *Ornithobacterium hominis* colonisation dynamics and interaction with the nasopharyngeal microbiome in a South African birth cohort"

<u>Supplementary Materials</u>

<u>Longitudinal context of *O. hominis* isolates</u>

Where 16S data was available, samples were targeted for culture based on proportional abundance of *O. hominis* to maximise the chances of success. As illustrated in Supplementary Figure S1, infants usually carried *O. hominis* for several months and the sample with the highest relative abundance for a given child was selected for culture.

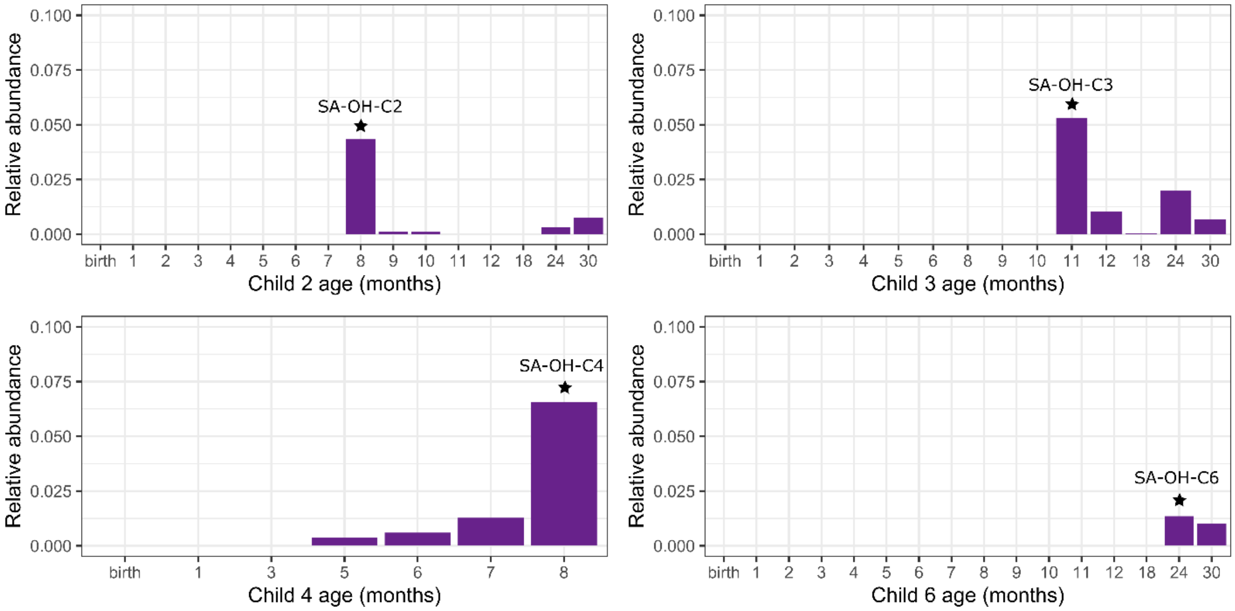


**Supplemental Figure S1: Proportional abundance of *O. hominis* reads in longitudinal samples from four children. Samples that were targeted for culture are indicated with a star and labelled with the isolate name.**

<u>16S rRNA gene analysis: selection of carrier and non-carrier groups</u>

The selection of 23 *O. hominis* carrier infants and 23 non-carriers from the screened dataset is illustrated in Figure S1. *O. hominis* carriers were infants who had at least one sample with a relative (raw read) abundance of *O. hominis* >1%. Non-carriers were infants who were never observed to carry *O. hominis* between the ages of 0 to 30 months, and for whom the longitudinal sampling density was at least 15 samples. Of the 100 infants who met these criteria, 23 were randomly assigned as pairs for the carrier group.

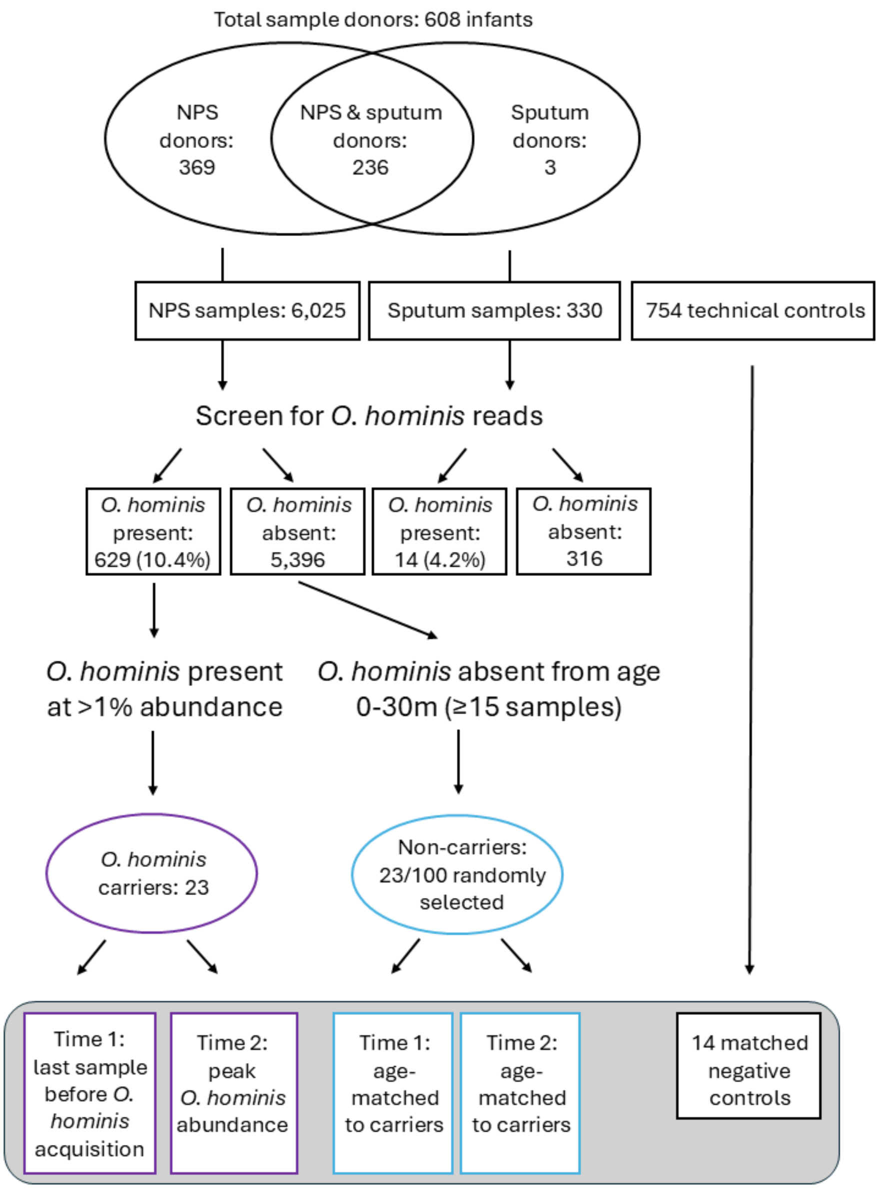


Figure S2: Flowchart of 16S rRNA data processing: from the full dataset of nasopharyngeal and sputum samples screened for *O. hominis* presence, to the subselection of *O. hominis* carriers and non-carriers used for nasopharyngeal microbial community analysis.

<u>Carrier and non-carrier covariates</u>

Information about the carrier and non-carrier groups is described in Table S1.

Table S1: Aggregated metadata for the *O. hominis* carrier and non-carrier groups

| | | **Carrier group** | **Non-carrier group** |
|---|---|---|---|
| Site | | | |
| | TC Newman | 10/23 (43.5%) | 10/23 (43.5%) |
| | Mbekweni | 13/23 (56.5%) | 13/23 (56.5%) |
| Birth season | | | |
| | Spring | 7/23 (30.4%) | 9/23 (39.1%) |
| | Summer | 5/23 (21.7%) | 8/23 (34.8%) |
| | Autumn | 3/23 (13.0%) | 4/23 (17.4%) |
| | Winter | 8/23 (34.8%) | 2/23 (8.7%) |
| Sex | | | |
| | Female | 14/23 (60.9%) | 13/23 (56.5%) |
| | Male | 9/23 (39.1%) | 10/23 (43.5%) |
| Gestational age at delivery (weeks) | | | |
| | Mean | 38 | 39 |
| | Median | 38 | 39 |
| | Min–Max | 33–41 | 32–42 |
| Birth weight (g) | | | |
| | Mean | 2895 | 3417 |
| | Median | 2900 | 3400 |
| | Min–Max | 1710–3940 | 2630–4440 |
| Delivery mode | | | |
| | Caesarean | 2/23 (8.7%) | 5/23 (21.7%) |
| | Vaginal | 21/23 (91.3%) | 18/23 (78.3%) |
| Maternal age at birth (years) | | | |
| | Mean | 26 | 28 |
| | Median | 24 | 29 |
| | Min–Max | 19–45 | 19–41 |
| Maternal HIV status | | | |
| | HIV negative | 18/23 (78.3%) | 20/23 (87.0%) |
| | HIV positive | 5/23 (21.7%) | 3/23 (13.0%) |
| Ethnicity | | | |
| | Black African | 13/23 (56.5%) | 13/23 (56.5%) |
| | Mixed ancestry | 10/23 (43.5%) | 10/23 (43.5%) |
| Maternal education | | | |
| | Lower than secondary | 18/23 (78.3%) | 15/23 (65.2%) |
| | Secondary or higher | 5/23 (21.7%) | 8/23 (34.8%) |
| Household income | | | |
| | <R1000/m | 10/23 (43.5%) | 9/23 (39.1%) |
| | R1000-5000/m | 12/23 (52.2%) | 12/23 (52.2%) |

| | | | |
|---|---|---|---|
| | >R5000/m | 1/23 (4.3%) | 2/23 (8.7%) |
| Parent employment | | | |
| | Not working | 17/23 (73.9%) | 12/23 (52.2%) |
| | Working | 6/23 (26.1%) | 11/23 (47.8%) |
| Marital status | | | |
| | Married/cohabiting | 5/23 (21.7%) | 10/23 (43.5%) |
| | Single | 18/23 (78.3%) | 13/23 (56.5%) |
| Number in household | | | |
| | Mean | 6 | 5 |
| | Median | 6 | 5 |
| | Min–Max | 1–12 | 2–10 |
| Prenatal smoking | | | |
| | No smoking | 14/23 (60.9%) | 20/23 (87.0%) |
| | Smoking | 9/23 (39.1%) | 3/23 (13.0%) |
| Prenatal alcohol exposure | | | |
| | Exposure | 5/23 (21.7%) | 1/23 (4.3%) |
| | No exposure | 18/23 (78.3%) | 20/23 (87.0%) |
| | Not recorded | 0 | 2/23 (8.7%) |
| Time breastfeeding (months) | | | |
| | Mean | 14 | 13 |
| | Median | 11 | 7 |
| | Min–Max | 0–26 | 0–26 |
| Time exclusive breastfeeding (months) | | | |
| | Mean | 3 | 2 |
| | Median | 2 | 1 |
| | Min–Max | 0–6 | 0–6 |

Samples from the *O. hominis* carrier and non-carrier groups were well matched for age, with a median difference in age of 2 days (maximum difference 37 days). The age distribution and gap between timepoints is illustrated in Figure S3.

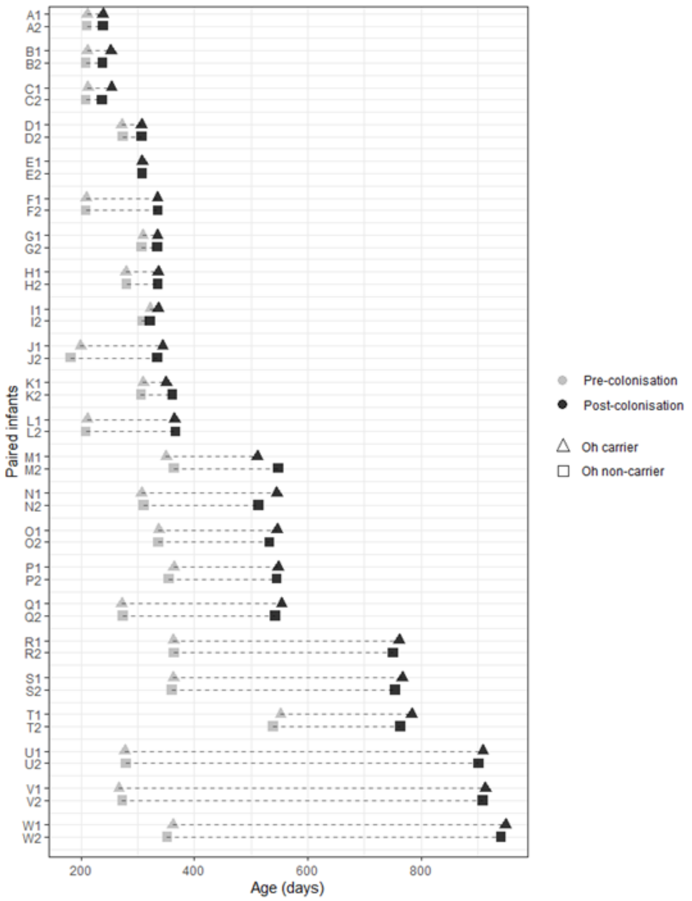


Figure S3: Age distribution of paired samples in the *O. hominis* carrier (triangle) and non-carrier (square) groups.

<u>Decontamination</u>

Following processing with Decontam, 350 ASVs were removed (comprising 2.3% of all reads including controls, but only 0.44% of reads from samples). Most removed taxa were plausible contaminants, such as *Anoxybacillus*, *Pelomonas*, *Methylobacterium*, and *Acinetobacter*. Others were human associated, notably *Cutibacterium* and *Staphylococcus*—skin microbes that are commonly reported contaminants in other studies (42). Although it may be appropriate to leave human-associated ASVs intact in some circumstances, these were removed in accordance with the Decontam results because the abundance in samples was extremely low while being almost ubiquitous in the negative controls, and so would not be included in downstream analysis (for example *Staphylococcus* mean abundance was 0.02% in samples but 7.5% in negative controls).

### Custom Refseq database

The reference database included all 99,432 genomes present in RefSeq from the following 31 genera: *Acinetobacter, Actinobacillus, Alloprevotella, Bergeyella, Campylobacter, Corynebacterium, Dolosigranulum, Filifactor, Fusobacterium, Gemella, Candidatus* Gracilibacteria*, Haemophilus, Helcococcus, Helicobacter, Johnsonella, Kocuria, Leptotrichia, Moraxella, Mycoplasma, Neisseria, Porphyromonas, Prevotella, Pseudomonas, Rappaport, Segatella, Staphylococcus, Streptobacillus, Streptococcus, Suttonella, Ureaplasma,* and *Veillonella*. Genes were reported that had >95% amino acid identity over >50% of their length.
